# Substrate channeling in oxylipin biosynthesis through a protein complex in the plastid envelope of *Arabidopsis thaliana*

**DOI:** 10.1101/286864

**Authors:** Stephan Pollmann, Armin Springer, Sachin Rustgi, Diter von Wettstein, ChulHee Kang, Christiane Reinbothe, Steffen Reinbothe

**Affiliations:** Centro de Biotecnología y Genómica de Plantas, Universidad Politécnica de Madrid (UPM) – Instituto Nacional de Investigación y Tecnología Agraria y Alimentación (INIA), Campus de Montegancedo, 28223 Pozuelo de Alarcón (Madrid), Spain; Medizinische Biologie und Elektronenmikroskopisches Zentrum (EMZ), Universitätsmedizin Rostock, D-18055 Rostock Germany; Clemson University, Department of Plant and Environmental Sciences, Pee Dee Research and Education Center, Florence, SC 29506, USA; Department of Crop and Soil Sciences, Washington State University, Pullman WA 99164, USA; Molecular Plant Sciences Program, Washington State University, Pullman WA 99164, USA; Center for Reproductive Biology, Washington State University, Pullman WA 99164, USA; Department of Chemistry, Washington State University, Pullman WA 99164, USA; School of Molecular Biosciences, Washington State University, Pullman WA 99164, USA; Biomolecular Crystallography Center, Washington State University, Pullman WA 99164, USA; Laboratoire de Bioénergétique Fondamentale et Appliquée, Université Grenoble Alpes, BP 53, Grenoble, Cedex F-38041, France

**Author notes:** Deceased April 13, 2017 (Diter von Wettstein). Correspondence or, Centro de Biotecnología y Genómica de Plantas (UPM-INIA). Campus de Montegancedo. Autopista M40 (km 38), 28223 Pozuelo de Alarcón, Madrid, Spain.

**Keywords:** Allene oxide synthase (AOS), Allene oxide cyclase (AOC), Chloroplast envelope protein complex, Hydroperoxide lyase (HPL), Lipoxygenase (LOX), Metabolite channeling, Plant defense

## Abstract

Oxygenated membrane fatty acid derivatives dubbed oxylipins play important roles in the plant’s defense against biotic and abiotic cues. Plants challenged by insect pests, for example, synthesize a blend of different defense compounds that, amongst others, comprise volatile aldehydes and jasmonic acid (JA). Because all oxylipins are derived from the same pathway, we asked how their synthesis might be regulated and focused on two closely related, atypical cytochrome P450 enzymes designated CYP74A and CYP74B, i.e., allene oxide synthase (AOS) and hydroperoxide lyase (HPL). Both enzymes compete for the same substrate but give rise to different products. While the final product of the AOS branch is JA, those of the HPL branch comprise volatile aldehydes and alcohols. AOS and HPL are plastid envelope enzymes in *Arabidopsis thaliana* but accumulate at different locations. Biochemical experiments identified AOS as constituent of complexes also containing lipoxygenase 2 (LOX2) and allene oxide cyclase (AOC), which catalyze consecutive steps in JA precursor biosynthesis, while excluding the concurrent HPL reaction. Based on published X-ray data, the structure of this complex could be modelled and amino acids involved in catalysis and subunit interactions identified. Genetic studies identified the microRNA 319 (miR319)-regulated clade of TCP (TEOSINTE BRANCHED/CYCLOIDEA/PCF) transcription factor genes and CORONATINE INSENSITIVE 1 (COI1) to control JA production through the AOS-LOX2-AOC2 complex. Together, our results define a molecular branch point in oxylipin biosynthesis that allows fine-tuning the plant’s defense machinery in response to biotic and abiotic stimuli.

## INTRODUCTION

Jasmonic acid (JA) and its derivatives are cyclopentanone compounds of widespread occurrence and ubiquitous function in plants (Böttcher and Pollmann, 2009; Reinbothe *et al*., 2009; Wasternack and Hause, 2013; Yan *et al*., 2013). JA biosynthesis comprises the release of linolenic acid from membrane lipids by phospho- and galactolipases, as well as 13-lipoxygenase (LOX), allene oxide synthase (AOS), and allene oxide cyclase (AOC) carrying out consecutive reactions in chloroplasts (Fig. 1). Whereas 13-LOX (EC 1.13.11.12) catalyzes the *regio*- and *stereo*specific hydroperoxidation of the C-13 atom of α-linolenic acid (α-LeA) giving rise to (13*S*)-hydroperoxylinolenic acid (13-HPOT), AOS (EC 4.2.1.92) converts 13-HPOT to 12,13-epoxylinolenic acid (EOT). Because EOT is short-lived and spontaneously disintegrates into volatile α- and γ-ketols as well as racemic 12-oxo-phytodienoic acid (OPDA), plants make use of AOC (EC 5.3.99.6) to assure *cis*-(+)-12-oxo-phytodienoic acid (*cis*-(+)-12-OPDA) synthesis. *cis*-(+)-12-OPDA then is exported from chloroplasts to the cytosol and transported further into peroxisomes, where the final reduction and ß-oxidation steps of JA biosynthesis take place. In *Arabidopsis thaliana*, six genes encode LOX isoforms, whereas one and four genes encode AOS and AOC enzymes, respectively. Mutant studies have provided valuable insights into the roles of the different LOX, AOS and AOC isoforms *in planta* (Schaller and Stintzi, 2009; Schaller *et al*., 2008; Wasternack and Hause, 2013; Yan *et al*., 2013).

**Figure 1.**
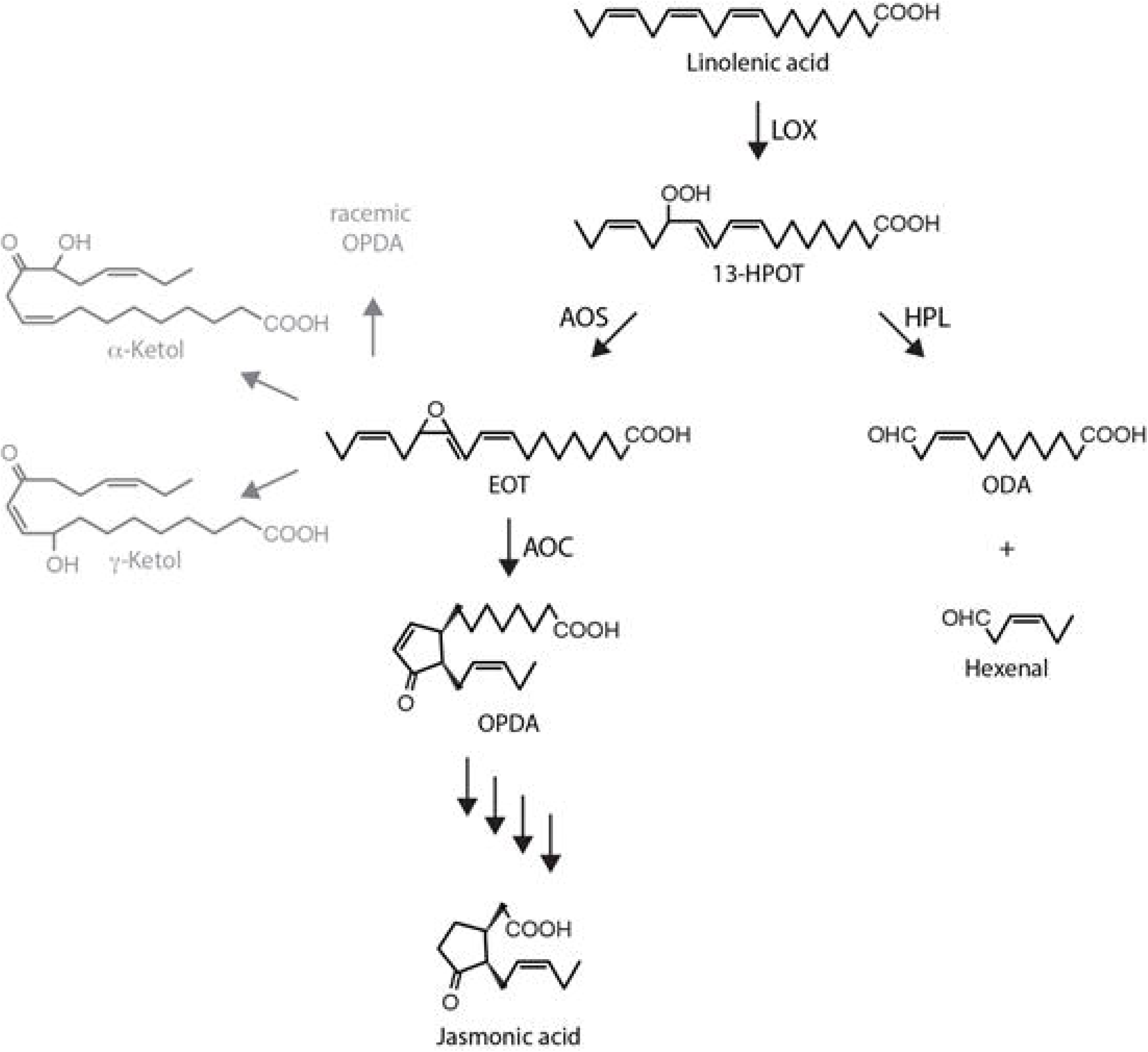
The Vick and Zimmerman pathway leading to JA and the concurrent AOS and HPL reactions. Pathway intermediates are abbreviated as follows: 13-HPOT, (9*Z*11*E*15*Z*13*S*)-13-hydroperoxy-9,11,15-octadecatrienoic acid; EOT, 12,13(*S*)-epoxy-9(*Z*),11,15(*Z*)-octadecatrienoic acid; OPDA, *cis*-(+)-12-oxophytodienoic acid; ODA, 12-oxo-*cis*-9-dodecenoic acid. The enzymes are indicated as follows: LOX, 13-lipoxygenase; AOS, allene oxide synthase; AOC, allene oxide cyclase; HPL, hydroperoxy lyase. Note that EOT is short-lived and spontaneously disintegrates into volatile α-ketols and γ-ketols as well as racemic OPDA.

(+)-*7-iso*-JA-Ile (JA-Ile) is the actual physiologically active compound (Fonseca *et al*., 2009). It triggers changes in gene expression including the activation of defense genes and inhibition of photosynthetic genes (Reinbothe *et al*., 1993a; Reinbothe *et al*., 1993b; Rustgi *et al*., 2014). Key regulatory elements in JA signaling involve the F-box protein COI1 (CORONATINE INSENSITIVE 1) acting as JA-Ile receptor (Xie *et al*., 1998; Yan *et al*., 2009), the E3 ubiquitin-ligase Skp-Cullin-F-box complex SCF^COI1^, and the JASMONATE ZIM-domain (JAZ) transcriptional repressors normally suppressing expression of JA response genes (Chini *et al*., 2007; Chung *et al*., 2008; Thines *et al*., 2007). Binding of JA-Ile to COI1 elicits the degradation of JAZ transcriptional repressors through the 26S proteasome and permits expression of JA response genes, driven by a number of MYC transcription factors (Chini *et al*., 2009; Fernández-Calvo *et al*., 2011; Hoffmann *et al*., 2011; Schweizer *et al*., 2013).

The Vick and Zimmerman pathway through which JA is produced has a number of branch points (Fig. 1). The question of how the flow of metabolites through the different branches is regulated in time and space is largely unanswered. In particular, several reactions compete for 13-HPOT *in planta* (Griffiths, 2015; Nilsson *et al*., 2016; Wasternack and Hause, 2013). One such reaction is catalyzed by fatty acid hydroperoxide lyase (HPL) that cleaves 13-HPOT into *Z*-3-hexenal and 12-oxo-*cis*-9-dodecenoic acid (ODA) of which c*is*-3-hexenal and the corresponding alcohol are volatile compounds operative in herbivore deterrence (Bate and Rothstein, 1998; Blée, 1998; Croft *et al*., 1993; Wu and Baldwin, 2010)(Fig. 1). Here, we report on the identification of a protein complex comprising LOX2, AOS and AOC2 in the plastid envelope of *Arabidopsis* chloroplasts that channels α-LeA into JA biosynthesis. Because the expression of LOX2, AOS and AOC2 is under the control of JA-responsive microRNAs and *COI1*, a mechanism is suggested that boosts JA over aldehyde production for rapid local and systemic defense gene activation in plants.

## RESULTS

### *In vitro*-import into chloroplasts and membrane targeting of AOS and HPL

AOS and HPL from *Arabidopsis thaliana* and other plant species belong to the family of atypical cytochrome P450s, designated CYP74. AOS (CYP74A) and HPL (CYP74B) require neither O_2_ nor NADPH-dependent cytochrome P450 reductase for activity and thus are non-canonical P450s (Froehlich *et al*., 2001; Laudert *et al*., 1996; Song and Brash, 1991). Previously described cDNAs for AtAOS and AtHPL (Froehlich *et al*., 2001; Laudert *et al*., 1996) were used for coupled *in vitro*-transcription/translation in wheat germ extracts. The ^35^S-methionine- or ^14^C-leucine-labeled proteins were then added to *Arabidopsis* chloroplasts that had been isolated by Percoll/sucrose density gradient centrifugation. Import experiments revealed that both AOS and HPL were taken up by isolated chloroplasts and processed by cleavage of their transit peptides (Fig. 2A). The resulting mature enzymes, however, were targeted to different locations (Fig. 2B-D). Whereas AOS was targeted to the inner envelope of chloroplasts, HPL was localized in the outer envelope fraction. Both enzymes were degraded by added trypsin, but only HPL was sensitive to thermolysin (Fig. 2A and B). Thermolysin is a protease that degrades surface-exposed proteins of the outer plastid envelope, whereas trypsin is known to access the intermembrane space between the outer and inner envelope, respectively (Cline *et al*., 1984; Kessler and Blobel, 1996). Thus, AOS is likely to face the intermembrane space as it was protected against thermolysin, whereas HPL is exposed to the cytosolic side of the outer plastid envelope and was, thus, sensitive to thermolysin. Western blot experiments revealed the co-localization of AOS with two translocon proteins of the inner chloroplast envelope, TIC55 and TIC110, and that of HPL with the translocon protein of the outer chloroplast envelope, TOC75 (Fig. 2C). Using TIC110 as diagnostic marker, any post-import degradation of ^35^S-AOS or ^35^S-HPL could be excluded (Fig. 2B, panel c). TIC110 is a very sensitive marker for monitoring the trypsin digestion procedure and easily undergoes degradation if no precautions are taken, such as inclusion of trypsin inhibitor during all chloroplast fractionation steps (Jackson *et al*., 1998). Extraction of outer and inner plastid envelopes with 1 M NaCl or 0.1 M Na_2_CO_3_, pH 11, revealed that HPL and AOS were tightly bound to their respective target membranes (Fig. 2D). The localizations of HPL and AOS confirmed previous data for chloroplasts from *Arabidopsis* (Joyard *et al*., 2010; Mwenda et al., 2015) and tomato (Froehlich *et al*., 2001), respectively, while not precluding divergent locations in other plant species.

**Figure 2.**
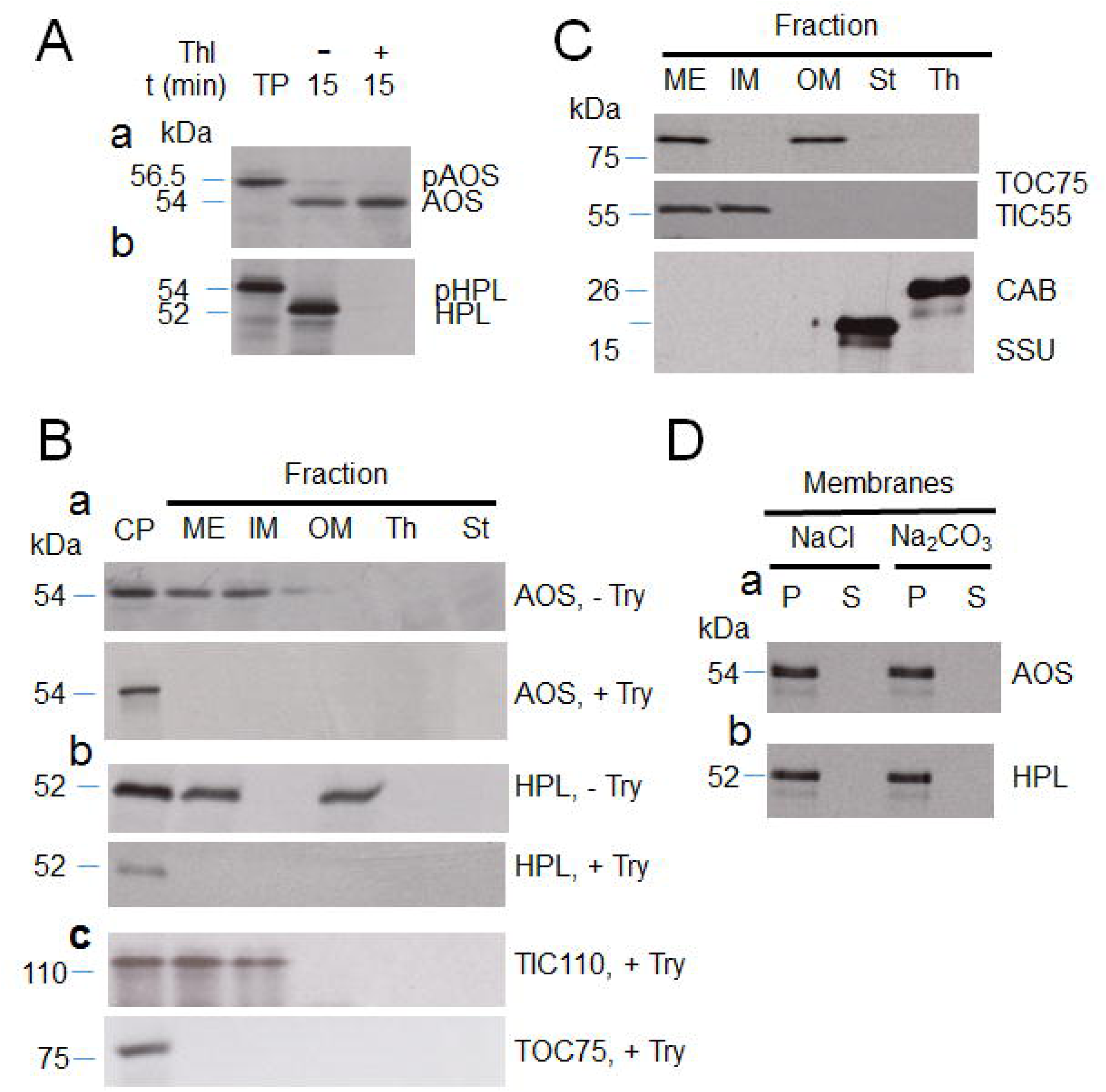
*In vitro*-import and differential membrane binding of AOS and HPL in chloroplasts. **(A)** Levels of ^35^S-Met-labelled (^35^S)-AOS (a) and ^35^S-HPL (b) before and after import into isolated *Arabidopsis* chloroplasts. Thl, thermolysin; TP, translation product. Positions of precursor protein (pAOS and pHPL) and mature protein (AOS, HPL) are indicated. **(B)** Detection by SDS-PAGE and autoradiography of ^35^S-AOS (a) and ^35^S-HPL (b) in trypsin (Try)-treated (+Try) and untreated (-Try) mixed outer and inner plastid envelopes (ME), inner plastid envelope (IM), outer plastid envelope (OM), thylakoids (Th) and stroma (St). CP defines the chloroplast reference fraction prior to import and protease treatment. The Western blot in panel c shows the levels of TIC110 and TOC75 in non-trypsin-treated chloroplasts (CP) versus trypsin (Try)-treated chloroplasts containing imported ^35^S-AOS/HPL and respective subfractions. **(C)** Western blot analysis of TOC75, the translocon at the inner chloroplast envelope membrane protein TIC55, the chlorophyll a/b binding protein LHCII (CAB) and the small subunit of ribulose-1,5-bisphosphate carboxylase/oxygenase (SSU) in the indicated fractions of non-trypsin-treated chloroplasts. Abbreviations are as in B. **(D)** Membrane binding of imported ^35^S-AOS (a) and ^35^S-HPL (b), as assessed by their extractability by 1 M NaCl and 0.1 M Na_2_CO_3_, pH 11. Both pellet (P) and supernatant (S) fractions, respectively, obtained after sedimentation of the membranes, were tested by SDS-PAGE and autoradiography for the two labeled proteins.

### Isolation of plastid envelope proteins interacting with AOS and HPL

We next attempted to identify proteins interacting with AOS in the envelope of *Arabidopsis* chloroplasts. COOH-terminal, hexa-histidine (His_6_)-tagged AOS (AOS-(His)_6_) was expressed in bacteria and purified to apparent homogeneity by Ni-NTA chromatography. The chemically pure protein was incubated with isolated chloroplasts in standard import reactions containing 2.5 mM Mg-ATP (Froehlich *et al*., 2001). After incubation, mixed outer and inner envelopes were isolated from ruptured chloroplasts (Li *et al*., 1991; Schnell *et al*., 1994) and solubilized with 1.3% decyl maltoside (Caliebe *et al*., 1997). Proteins adhering to AOS-(His)_6_ were subjected to size exclusion chromatography and AOS-(His)_6_ detected with AOS-specific or (His)_6_ antibodies (Froehlich *et al*., 2001; Laudert *et al*., 1996). For comparison, AOS-containing plastid envelope complexes were isolated from transgenic plants overexpressing Flag-tagged AOS (AOS–Flag) under the control of the 35S cauliflower mosaic virus promoter.

Figure 3A and B depict the elution profiles of AOS after its purification from either isolated chloroplasts following the *in vitro*-import reaction of AOS- (His)_6_ or transgenic plants overexpressing Flag-tagged AOS. Whereas AOS-Flag and AOS-(His)_6_ were present in higher molecular mass complexes of ≈250 kDa (peak I), a second, lower molecular mass complex was only detectable for AOS-Flag (peak II). The lack of a corresponding AOS-(His)_6_ peak may be due to the lower abundance of this protein in the *in vitro*-uptake assays with isolated chloroplasts. When the total pattern of proteins that co-purified with AOS-(His)_6_ in peak I was analyzed by SDS-PAGE, at least 10 bands were found (Fig. 3C, lane 1) of which, besides the bait protein, the ≈100 kDa band was identified by protein sequencing as LOX2 and the ≈72 kDa and ≈24 kDa bands as AOC2, forming SDS-resistant trimers and monomers, respectively. When similar experiments were carried out with HPL-(His)_6_, a completely different protein pattern was obtained which, most notably, did not contain any of the proteins detected with AOS-(His)_6_. (Fig. 3C, lane 2).

**Figure 3.**
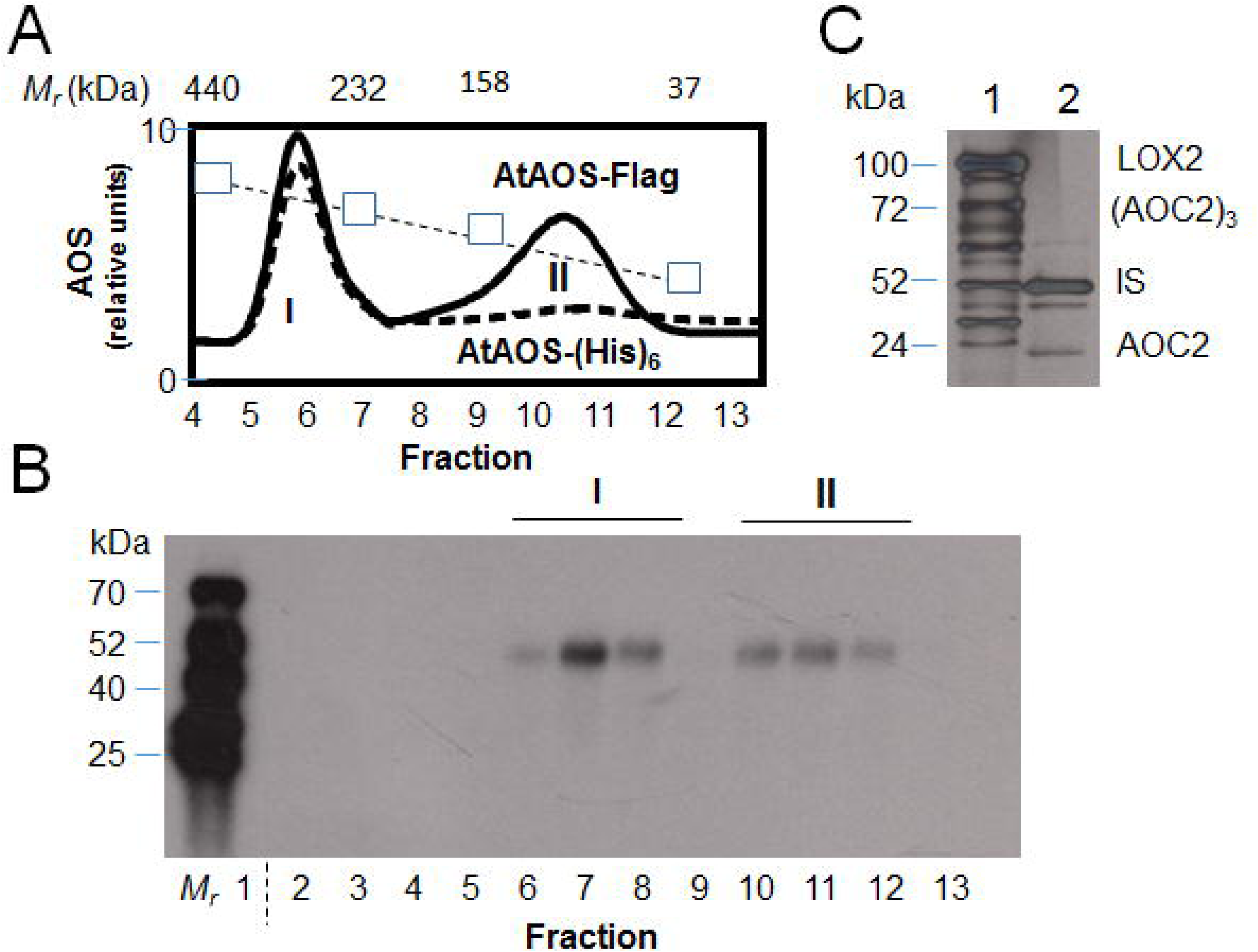
Detection of AOS complexes *in vitro* and *in planta*. **(A)** Gel filtration elution profile of AOS-Flag in protein extracts of transgenic plants expressing Flag-tagged AOS (solid line) and in isolated chloroplasts after *in vitro*-import of AOS-(His)_6_ (broken line). Positions of size marker proteins are indicated (squares and dotted line). **(B)** as A, but showing a Western blot analysis of AOS-Flag. *M_r_* defines radioactive molecular mass standards. **(C)** SDS-PAGE pattern of plastid envelope proteins co-purifying with AOS-(His)_6_ (lane 1) and HPL-(His)_6_ (lane 2). The indicated bands were identified by protein sequencing.

### Genetic evidence for the existence of a chloroplast envelope complex comprising AOS, LOX2 and AOC2

Because the expression of *LOX2*, *AOS* and *AOC2* in *Arabidopsis* is under the control of the microRNA 319 (miR319)-regulated clade of *TCP* (*TEOSINTE BRANCHED/CYCLOIDEA/PCF*) transcription factor genes (Schommer *et al*., 2008), we used the *jaw-*D mutant with strongly reduced mi319-dependent expression of JA biosynthetic genes to investigate the interaction of LOX2, AOS and AOC2 genetically. When *jaw-*D plants, in which *TCP4* expression is strongly down-regulated, and primary transformants expressing a miR319-resistant version of *TCP4* (referred to as *TCP4* plants in the following) that dominantly regulates *LOX2* expression (Palatnik *et al*., 2003; Schommer *et al*., 2008), were grown for 14 d in a greenhouse, not much of a difference in pheno-type was seen. However, this picture changed at later stages of plant development and especially at the stage when plants matured and entered senescence. In agreement with previous results (Schommer *et al*., 2008), *TCP4* plants then displayed a marked acceleration of senescence, whereas *jaw-*D plants exhibited a tremendous, 2 week-delay in leaf senescence. These differential changes correlated with altered patterns of plastid envelope proteins comprising LOX2, AOS and AOC2 (Fig. 4; see also Supplemental Information file I showing respective densitometric analyses using the NIH endorsed ImageJ software, https://imagej.net/). *TCP4* plants more rapidly accumulated all three JA biosynthesis enzymes than wild-type plants (Fig. 4A)

**Figure 4.**
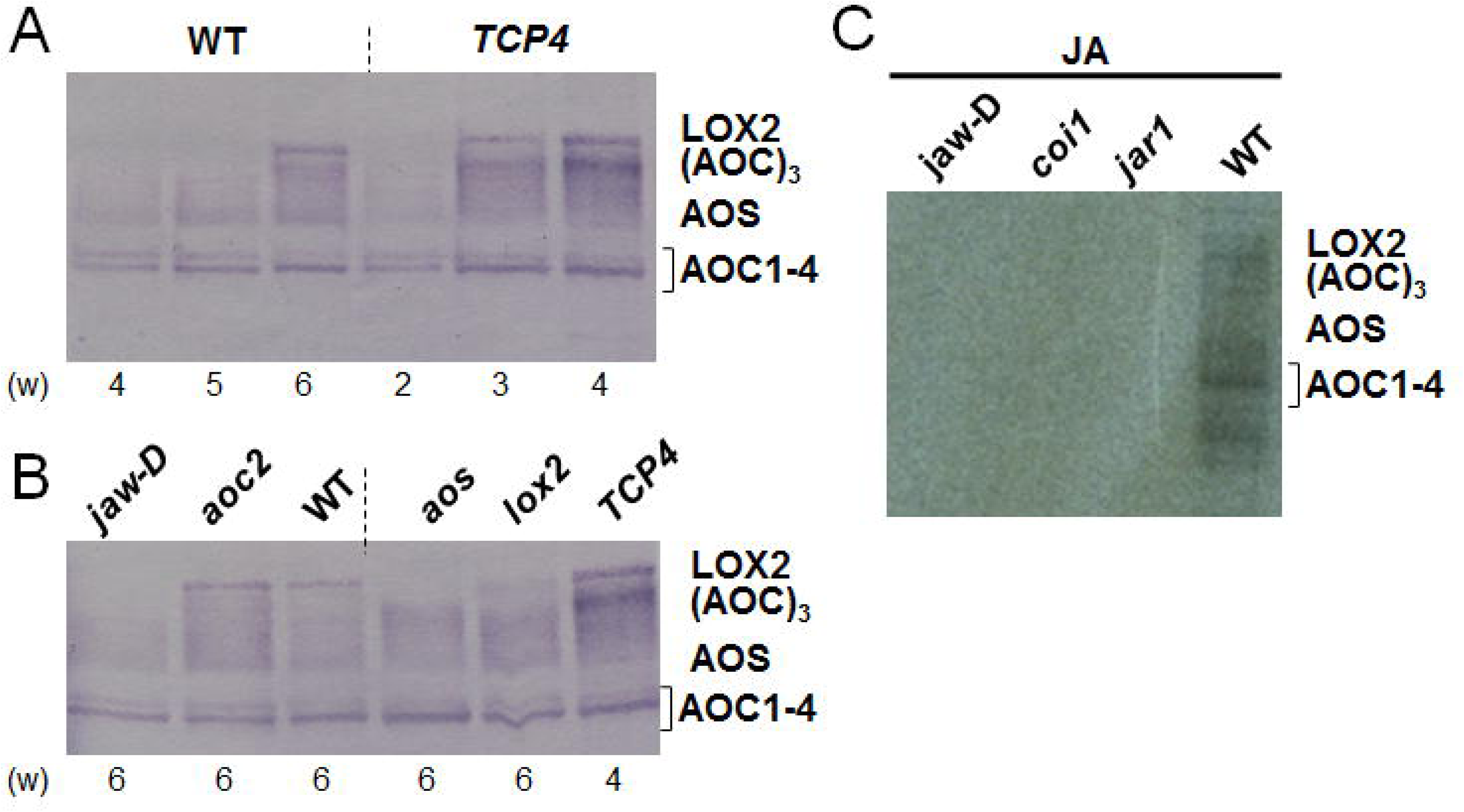
Genetic dissection of LOX2-AOS-AOC2 complex formation. **(A)** Time course of AOS, LOX2 and AOC2 accumulation in wild-type (WT) and *TCP4* plants expressing a mi319-resistant version of TCP4 during development (in weeks [w]). **(B)** as A, but depicting accumulation of AOS, LOX2 and AOC in plants of *jaw-*D, *aoc2*, wild-type (WT), *lox2*, *aos* and *TCP4* backgrounds. **(C)** as A, but showing accumulation of AOS, LOX2 and AOC in plastid envelopes of 4 weeks-old *jaw-*D, *coi1*, *jar1* and wild-type (WT) plants after treatment with 45 µM MeJA for 24 h. Western blots were simultaneously probed with antisera against LOX2, AOS and AOC2 and developed with either horseradish peroxidase-alkaline phosphatase-based (A and B) or enhanced chemiluminescence-based (C) detection systems.

JA biosynthesis is under the control of a feed-forward loop and consequently can be boosted by exogenously added JA (Wasternack and Hause, 2013; Yan *et al*., 2013). To pinpoint the role of LOX2, AOS and AOC2 in this loop, JA-deficient seedlings of the *aos* mutant (Park *et al*., 2002; von Malek *et al*., 2002) and JA-Ile-insensitive seedlings of the *coi1* mutant (Feys *et al*., 1994) were used. For comparison, we employed *Arabidopsis* mutants defective in the *LOX2* and *AOC2* gene, respectively (Seltmann *et al*., 2010; Stenzel *et al*., 2012). When the patterns of plastid envelope proteins were compared, some differential effects were observed. As displayed in Fig. 4B, the *aoc2* knock-out mutants contained wild-type levels of AOC1, AOC3 and AOC4 as well as wildtype levels of *AOS* and *LOX2*. By contrast, *aos* and *lox2* mutants likewise expressed reduced amounts of LOX2 and AOC2 trimers, while retaining the levels of AOC1-4 monomers. These findings were suggestive of partially overlapping roles and expression patterns of the four AOC family members (see Supplemental Information file II; Fig. S1-3), with AOC1, AOC3 and AOC4 presumably being capable of replacing AOC2 forming homo- and hetero-complexes. It has recently been demonstrated that all four AOCs display identical catalytic activities (Otto *et al*., 2016).

To further dissect the feed-forward loop operating in JA biosynthesis, seedlings of the JA-insensitive *coi1* mutant (Feys *et al*., 1994) and JA-resistant *jar1* mutant (Staswick *et al*., 1992) were used. *JAR1* encodes an enzyme that provides the physiologically active JA-Ile conjugate for CORONATINE INSENSITIVE 1 (COI1)-dependent signal transduction (Feys *et al*., 1994; Staswick *et al*., 1992). *coi1* and *jar1* seedlings were grown for 14 days in continuous white light and in turn sprayed with MeJA. MeJA treatment triggered expression of LOX2, AOS and AOC1-4 in wild-type plants but not in *jaw*-D, *coi1* and *jar1* plants (Fig. 4C). Hence, JA production was required and relied on an intact JA signal transduction chain comprising JA-Ile and COI1, linking JA signal with miR319-regulated TCPs.

### Probing the interaction of LOX2, AOS and AOC2 through crosslinking

The presence of higher molecular mass complexes containing LOX2, AOS and AOC2 in the envelope of *Arabidopsis* chloroplasts suggested the possibility that the enzymes may physically and functionally interact in JA precursor biosynthesis. As a first step to test this hypothesis, crosslinking experiments were conducted using bacterially expressed, (His)_6_-tagged, purified precursors that had been derivatized with ^125^I-N-4[(p-azidosalicyl-amido)butyl]-3’(2-pyridyl-dithio)propion-amide (**^125^**I-APDP), which is a hetero-bifunctional, photoactivatable and cleavable label-transfer crosslinker frequently used in the chloroplast protein import field (e.g., Ma *et al*., 1996). ^125^I-APDP contains a 21 Å spacer arm that, depending on its location on the bait protein, can penetrate to different extents into the respective target membrane and label proteins and lipids.

^125^I-APDP-labeled LOX2, AOS and AOC2, respectively, were imported in darkness into isolated *Arabidopsis* chloroplasts (Springer et al, 2016; Figs. 2 and S4). After 15 min incubation, label-transfer crosslinking was induced by UV-light exposure on ice. Envelopes, in turn, were isolated from ruptured chloroplasts, solubilized with 3% SDS (Ma *et al*., 1996), and ^125^I-APDP-labeled proteins detected by SDS-PAGE and autoradiography. Figure 5A revealed different banding patterns for the three labeled proteins. LOX2 and AOC2 (being present as a trimer), both interacted with AOS, with little or no direct interaction between the two of them. For ^125^I-APDP-AOS, label transfer occurred onto LOX2 and AOC2 (Fig. 5A; see also the densitometric scans in Supplemental Information file I). Interestingly, ^125^I-APDP-AOS gave rise to both, ^125^I-labeled AOC2 trimers and monomers (Fig. 5A). AOC2 monomers were also seen in assay mixtures containing ^125^I-APDP-LOX2. This observation could be suggestive of a shuffling of monomers and trimers of AOC2 into the envelope complex. On the other hand, the nature of the chosen crosslinker and its topology on the bait protein may explain this result. Last but not least, it is also possible that substrate binding and conversion affected the interaction of LOX2, AOS and AOC2 and thereby had an impact on the crosslinking results.

**Figure 5.**
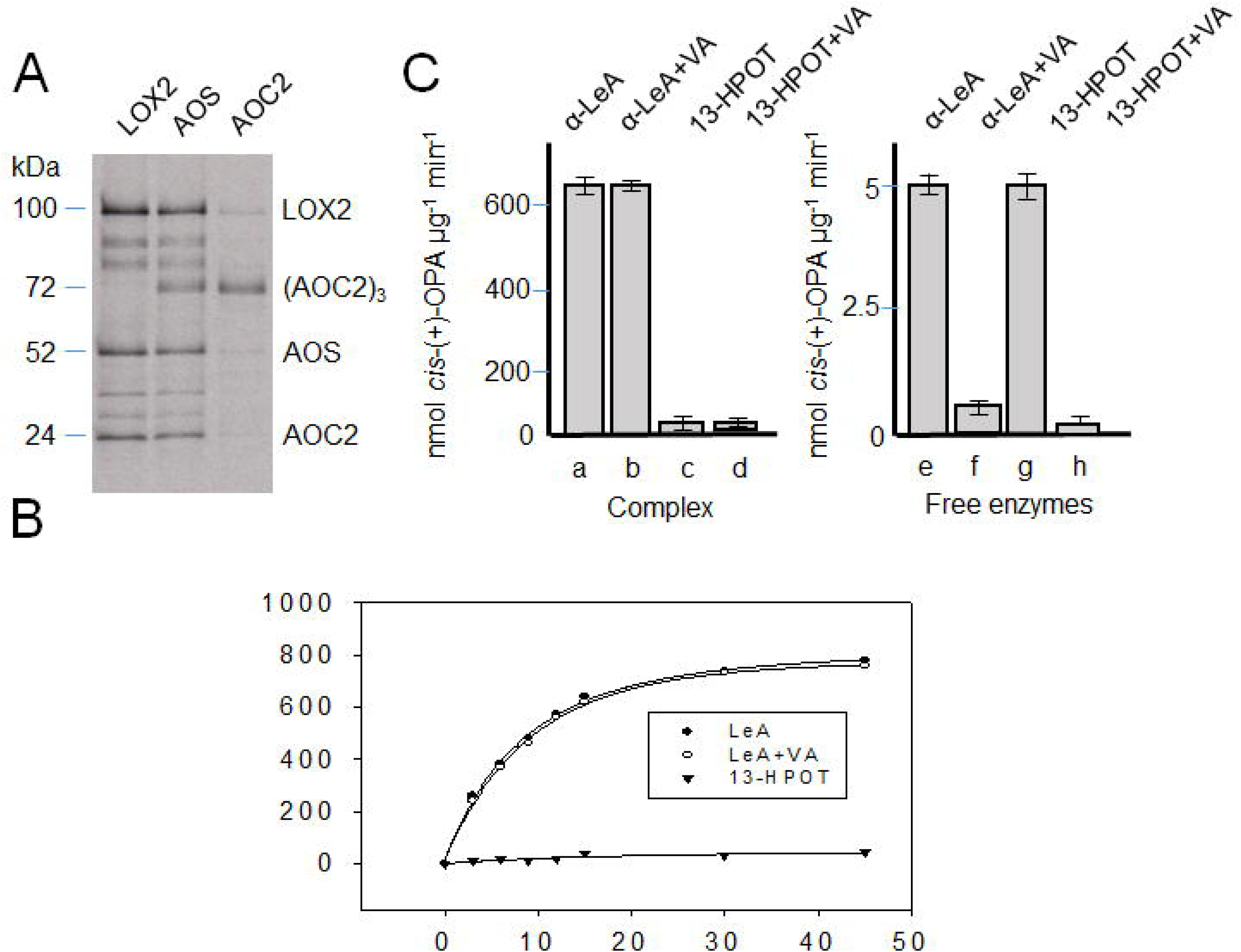
Channeling of jasmonate precursors in the LOX2-AOS-AOC2 plastid envelope complex. **(A)** Label-transfer crosslinking by ^125^I-APDP–LOX2, ^125^I-APDP–AOS and ^125^I-APDP–AOC2 of proteins in mixed outer and inner envelopes of *Arabidopsis* chloroplasts. **(B)** Time course of *cis*-(+)-12-OPDA formation from α-LeA (filled circles) and 13-HPOT (filled triangles) by the isolated envelope complex containing LOX2, AOS and AOC2. For comparison, *cis*-(+)-12-OPDA formation from α-LeA was tested in the presence of vernolic acid (VA)(open circles). **(C)** Rate of *cis*-(+)-12-OPDA formation determined after 10 min incubations for α-LeA (lanes a and e) and 13-HPOT (lanes c and g) by virtue of the LOX2-AOS-AOC2 envelope complex (lanes a-d), as compared to free enzymes released from the membrane complex by treatment with 3% SDS and dialyzed before analysis (lanes e-h, respectively). Respective controls show incubations performed with vernolic acid (VA)(lanes b, d and f and h, respectively). Error bars refer to three independent experiments.

With respect to the overlapping expression pattern of the *AOC* isogenes (see Supplemental Information file II; Figs. S2D and S3), we additionally asked whether the respective isoenzymes are capable of forming heteromeric complexes within the chloroplast, or whether complex formation is restricted to the formation of homomers. By using a bimolecular fluorescence complementation (BiFC) approach, it was possible to demonstrate that there are no restrictions in complex formation, as all possible interactions of AOC1–4 have been detected by fluorophore complementation in chloroplasts of *Arabidopsis* protoplasts (Fig. S2). These results are in agreement with recent data by Otto *et al*. (2016) who studied the capabilities of all four AOC isoforms to form trimers and how this may affect enzyme activity.

If LOX2, AOS and AOC2 were to structurally and functionally interact, they could provide the possibility of substrate channeling without concurrent side reactions in *cis*-(+)-12-OPDA synthesis from α-LeA. To address this possibility, bacterially expressed and purified, non-^125^I-APDP-derivatized AOS-(His)_6_ was imported into *Arabidopsis* chloroplasts. Then, the plastids were lysed and protein complexes containing AOS-(His)_6_ isolated by affinity chromatography from 1.3% decyl maltoside-solubilized envelopes. For reference, SDS-dissociated and dialyzed envelope complexes containing the free LOX2, AOS and AOC2 in amounts identical to those in the intact complexes were used. Activity measurements using α-LeA and 13-HPOT revealed a tight channeling of metabolites in the intact envelope complex. This is apparent from time courses of *cis*-(+)-12-OPDA formation from α-LeA and 13-HPOT (Fig. 5B) and respective quantification of *cis*-(+)-12-OPDA (Fig. 5C). While α-LeA was converted to *cis*- (+)-12-OPDA by the isolated, intact (non-SDS-dissociated) plastid envelope complex, 13-HPOT was not accepted as substrate. In line with the tight channeling of metabolites, the AOC inhibitor vernolic acid ((+/-)-cis-12,13-epoxy-9(Z)-octadecenoic acid) (Hofmann *et al*., 2006) was unable to impede *cis*-(+)-12-OPDA synthesis (Fig. 5C). When applied to the SDS-dissociated, dialyzed complex containing free LOX2, AOS and AOC2, however, vernolic acid did block *cis*-(+)-12-OPDA synthesis from α-LeA and 13-HPOT (Fig. 5C). Remarkably, the rate of *cis*-(+)-12-OPDA formation from α-LeA in the intact complex was 120-fold higher than that in the free-enzyme assay (Fig. 5C), suggesting tight functional LOX2-AOS-AOC2 interactions to occur and to boost *cis*-(+)-12-OPDA synthesis. In summary, we obtained 2-fold higher activities for the free-enzyme assay as compared to Otto *et al*. (2016), i.e., 5 nmoles OPDA per min^−1^ µg^−1^ AOC2 and a 240-fold greater activity for the complex-containing assay (600 nmoles OPDA per min^−1^ µg^−1^ AOC2).

### Interaction of LOX2, AOS and AOC2 probed in the Split-Ubiquitin system

As an independent approach to confirm the interaction of LOX2, AOS and AOC2, we carried out split-ubiquitin yeast two-hybrid screens for membrane bound proteins. Yeast cells (strain DSY1) were first transformed with AOS- and LOX2-containing bait-vectors (AOS-Cub, LOX2-Cub), respectively, which express the bait proteins fused to the C-terminal part of UBIQUITIN. For each bait protein, we constructed and tested three different vectors (pAMBV4, pCMBV4, pTMBV4), providing promoters of different strength to drive bait protein expression. In our hands, the pTMBV4 vector that contains a highly potent TEF1 promoter gave the clearest results. Co-transformation was accomplished in a second, separate step in which AOS- and AOC2-containing prey vectors were introduced into the selected yeast cells (NubG-AOS, NubG-AOC2). As prey vectors, we used constructs that added the N-terminal part of UBIQUITIN to the N-terminal extremity of the preys (pADSL-Nx). In addition to the specific constructs used to study potential LOX2-AOS-AOC2-interactions, we used control vectors provided with the system to test for unspecific and auto-activation, respectively (NubG-Alg5), as well as positive (Alg5-Cub, NubI-Alg5) and negative (Alg5-Cub, NubG-Alg5) system controls. The co-transformed yeast cells were subjected to qualitative His-complementation growth tests carried out on SD plates supplemented with 5 mM 3-AT (3-amino-1,2,4-triazole) lacking the three amino acids Leu, Trp, and His. Additionally, the relative ß-galactosidase activities of the investigated co-transformed yeast strains were determined to provide quantitative data on the interaction strength of the analyzed protein pairs. According to the results summarized in Fig. 6, interactions were observed between AOS and AOC2, with a fraction of AOS presumably forming dimers, as found for AOS from *Parthenium argentatum* (Chang *et al*., 2008; Li *et al*., 2008). Some interactions could also be detected for LOX2 with AOC2 and AOS, although these were quite variable and dependent on the respective bait and prey protein.

**Figure 6.**
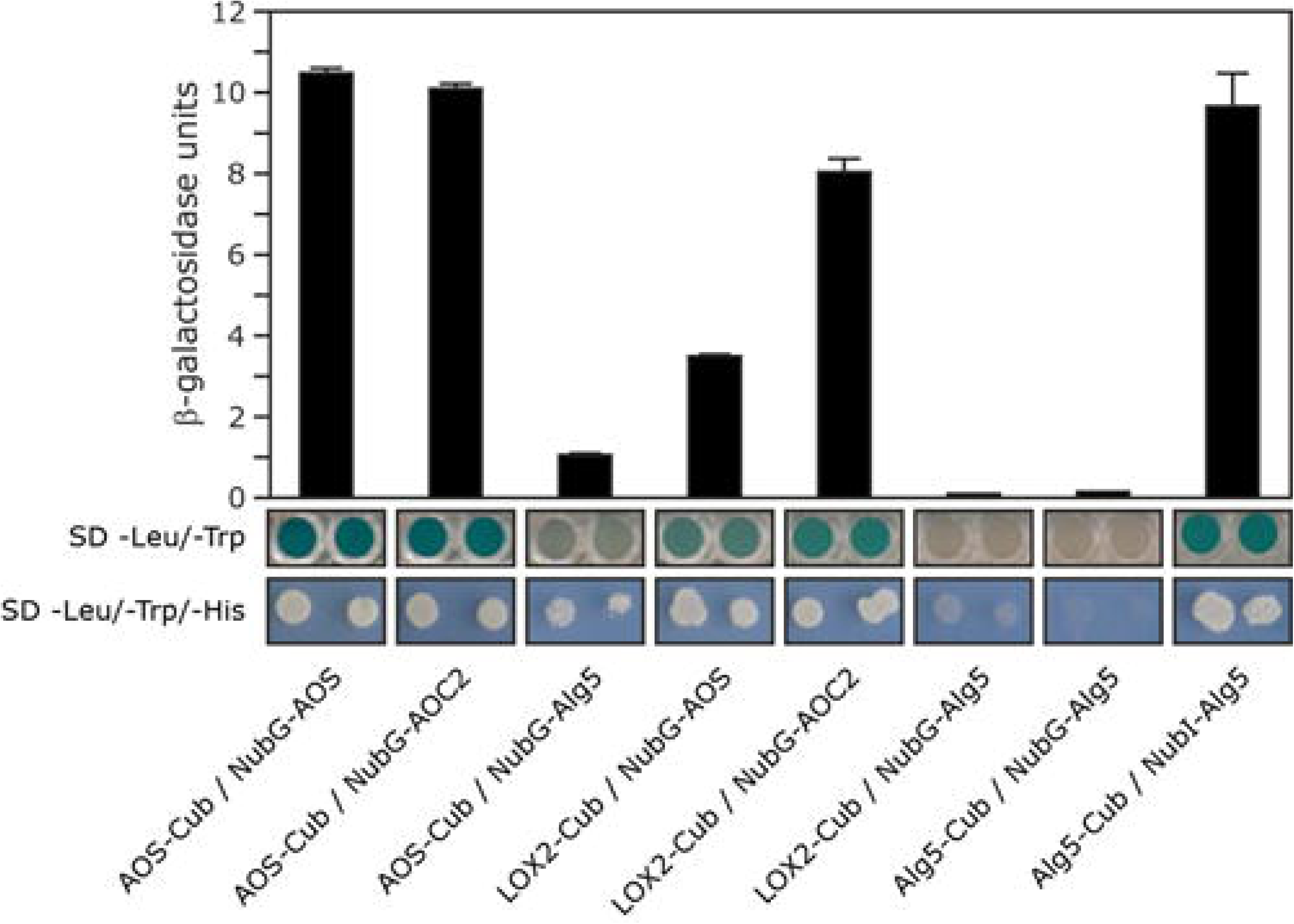
Genetic dissection of AOS-LOX2-AOX2 interactions in the split-ubiquitin system. Experiments were performed employing fusions to the C- (Cub) and N-terminal (NubG) halves of ubiquitin. Alg5-NubI, the fusion of the unrelated ER membrane protein Alg5 to NubI, was used as positive control. Negative controls were fusions of Alg5 to Cub or NubG (Alg5-Cub and Alg5-NubG). Qualitative estimation of interactions was deduced from the growth behavior of co-transformed yeast cells on solidified selection medium, while for quantitative assays the ß-galactosidase activity of cells grown in liquid culture was analyzed.

## DISCUSSION

### Sequestration of jasmonate precursor biosynthesis through a protein complex in the plastid envelope

In the present study, evidence is provided for the existence of a protein complex involved in JA precursor biosynthesis in *Arabidopsis* chloroplasts. We show that LOX2, AOS and AOC2, the enzymes that catalyze consecutive steps in JA precursor biosynthesis (Fig. 1), are co-localized in the inner envelope of *Arabidopsis* chloroplasts (Figs. 2 and S4; see also Springer et al., 2016), and formed complexes operating in *cis-*(+)-12-OPDA synthesis from α-LeA (Figs. 3, 5 and S5). It was possible to reconstitute a similar complex *in vitro* from soluble LOX2, AOS and AOC2 enzymes and isolated plastid envelope lipids (Fig. S5). In either case, LOX2, AOS and AOC2 were present in a 1:1:4 stoichiometry, indicating that these complexes may contain both, AOC2 monomers and trimers. The significance of this observation and the possibility of activity regulation through hetero-trimerization of AOC2 with AOC1, 3 and 4 remains to be established. It must be noted in this context, however, that also Zerbe and co-workers used AOS and AOC2 at a 1:4 molar ratio (Zerbe *et al*., 2007). Interaction studies in the split-ubiquitin yeast two-hybrid system confirmed the interaction of AOS and AOC2 (Fig. 6). When isolated plastid envelope complexes were supplied with α-LeA, only *cis*-(+)-12-OPDA accumulated in the reaction medium (Fig. 5). No evidence was obtained for the release of significant amounts of HPOT or EOT and its short-lived disintegration products (α-ketols and γ-ketols) as well as of racemic OPDA into the reaction mixture. These findings are suggestive of a tight channeling of metabolites through both, the isolated and reconstituted protein complexes. By virtue of the observed channeling of metabolites, the dilution of the reaction intermediates was kept low. On the other hand, the observed channeling of metabolites tremendously increased the rate of formation and, thus, yield of *cis-*(+)-12-OPDA from α-LeA. Compared to the free-enzyme assay, 120-fold higher activities were measured in the isolated envelope complex. Our data confirm and extend previous findings by Zerbe and colleagues who reconstituted *cis-*(+)-12-OPDA synthesis from 13-HPOT by combining purified recombinant AOS and AOC2 *in vitro* (Zerbe *et al*., 2007). The authors found that both soluble and matrix-bound enzymes are active and that their co-fixation on a solid matrix increased the yield of *cis-*(+)-OPDA from 13-HPOT by about 50%. In contrast to these studies, however, in which AOS and AOC2 were randomly bound to the matrix, thus excluding tight substrate channeling, in our experiments the enzymes were orderly associated, thereby allowing the channeling of α-LeA. Because vernolic acid failed to inhibit *cis-*(+)-12-OPDA synthesis from α-LeA or 13-HPOT in the native and reconstituted complexes (Fig. 5, Fig. S5), we conclude that the active site of AOC2 is largely inaccessible to the inhibitor. On the other hand, exogenously administered 13-HPOT was not accepted as substrates for *cis-*(+)-12-OPDA synthesis by the native and reconstituted complexes, but it was accepted in case of the free-enzyme assay (Fig. 5C). On the basis of these results we conclude that *cis-*(+)-12-OPDA synthesis from α-LeA is strictly compartmentalized, presumably to prevent competing side reactions such as that from 13-HPOT catalyzed by HPL or by chemical decay, which give rise to α- and γ-ketols. By this means, plants avoid the costly formation of *n*-hexenal and 12-oxo acids implicated in direct and indirect defenses of herbivores (Hoffmann *et al*., 2011; Wu and Baldwin, 2010), while maintaining the capacity to respond to biotic foes and abiotic stresses (Wasternack and Hause, 2013; Yan *et al*., 2013).

### Structural modeling of the LOX2-AOS-AOC2 envelope complex

We conclude from our results that AOS forms complexes with both LOX2 and AOC2 *in planta* as well as *in vitro*. Molecular modeling was carried out to obtain insight into the potential structure of this complex, using published X-ray structure data for soybean lipoxygenase L3 (PDB ID 1LNH;; Skrzypczak-Jankun *et al*., 1997; Youn et al., 2006), AOS (D 3CLI; Lee *et al*., 2008) and AOC2 (PDB ID 2GIN; Hofmann *et al*., 2006), obtained at 2.60 Å, 1.80 Å and 1.80 Å resolution, respectively, as templates (Figs. S6-S8). First, homology modelling was done for LOX2, based on the X-ray structure of soybean LOX L3 (Cho and Stahelin, 2006; Youn et al., 2006). The established LOX2 structure of Arabidopsis (Fig. S6) was then used in modelling of the whole, LOX2-, AOS- and AOC2-containing complex. The top rank model depicted in Fig. 7 suggests that at least two amino acid residues, Ser92 and Gly94, of LOX2 are potentially involved in the LOX2-AOS interaction, forming hydrogen bonds with Ser272 of AOS (Fig. S9A). The interacting Ser and Gly residues of LOX2 are not conserved between LOX2 and soybean LOX3 and reside in a loop region between β-sheets 1 and 2 of LOX2. Similarly, one region of interacting amino acids could be identified for each of the hypothetical LOX2-AOC2 and AOS-AOC2 complexes. In these complexes, Asp96 of LOX2 is predicted to interact with Asn42 of AOC2, and Phe44 and Ser45 of AOC2 were found when analyzing these enzymes apart from LOX2, while not seeing them in the modeled LOX2-AOS-AOC2 complex (Fig. S9B). Along with the crosslinking data, these findings are suggestive of a dynamic equilibrium of binding and release of all three enzymes to and from the complex.

**Figure 7.**
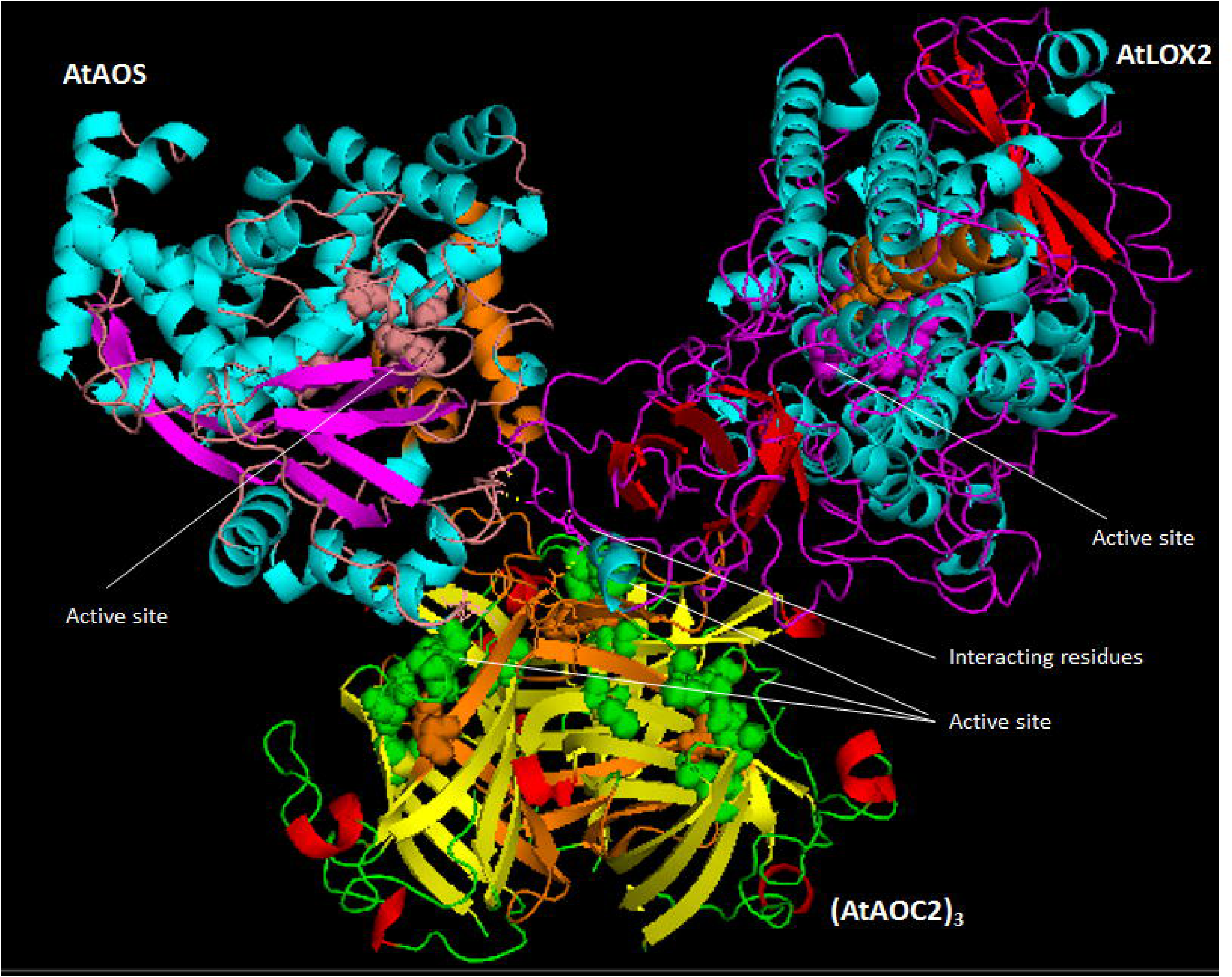
Structural model of the LOX2-AOS-(AOC2)_3_ interaction in the chloroplast inner envelope. Trans-membrane domain was shown in orange. Active site residues are shown by shears, and interacting residues by line. Active site predictions for LOX2 are based on Youn et al. (2006), for AOS2 on Lee et al. (2008) and for AOC2 on Hofmann et al. (2006) and trans-membrane domain predications are based on TMpred for LOX2 and (AOC2)_3_.

Details of how the LOX2-AOS-AOC2 complex may be bound to the membrane remain elusive. The fact that a functional complex could reconstituted from soluble LOX2, AOS and AOC2 enzyme molecules at first glance suggests that the lipid bilayers might be dispensable for formation of the whole complex. Nevertheless, all three enzymes attained salt- and protease-resistant states after respective *in vitro*-import reactions with chloroplasts, suggesting their tight binding to the lipid bilayers of the inner plastid envelope. Hydrophobic transmembrane (TM) domains may anchor LOX2, AOS and AOC2 in the membrane. Lee et al. (2008) identified several non-polar detergent binding α-helices in AOS of which those designated α-F and α-G were tentatively defined as putative TM domains in our structural models (Fig. S7A). Predictions made with TMpredn however, suggest a different part of the AOS polypeptide to form a single TM domain and that this overlaps with some of the active site residues (Fig. S7B). Such overlap would resemble that in the AOC2 trimer where some active site residues appear to be embedded into/are part of the predicted TM domain (Fig. S8). However, it cannot be excluded that some of the hydrophobic β-sheets forming the characteristic beta-barrel cavity are involved in membrane binding (Fig. S8). Lastly, the LOX structure of Skrzypczak-Jankun et al. (1997) used for modelling LOX2 from *Arabidopsis* (Fig. S6) suggests a hydrophobic environment of the catalytic pocket that could overlap with the predicted TM domain. Further work is needed to resolve the structure of the LOX2-AOS-AOC2 complex and elucidate its membrane binding.

It is attractive to hypothesize that LOX2 and AOS compete for binding sites on AOC2, thereby provoking the formation of AOC trimers, possibly involving other available AOC isoforms, too (Hofmann and Pollmann, 2008; Hofmann *et al*., 2006; Otto *et al*., 2016). As described by Otto *et al*. (2016), trimerization of AOC isoenzymes is likely to contribute to activity regulation. Several salt bridges between monomers and a hydrophobic core within the AOC2 trimer were identified and functionally proven by site-directed mutagenesis. While Lys152 of one monomer and Glu128 of the neighboring monomer established the observed salt bridges, amino acids involved in building the hydrophobic core of the trimer comprised Leu40, Leu50, Leu53 and Ile79 of all three monomers (Otto *et al*., 2016). Notable are the conservation of interacting amino acids and the overlapping expression patterns of AOC1/2 during germination, leaf production, rosette growth, inflorescence emergence and flowering with each other and with those of LOX2 and AOS (Fig. S1-3). By contrast, the expression pattern of AOC3 and AOC4 is more distinct (Fig. S1-3). On the other hand, HPL expression is comparably low in all of the developmental stages analyzed (Fig. S3). Together, these correlative data are suggestive of a co-evolution of mechanisms that allow favoring JA precursor biosynthesis over volatile production during plant development. This situation obviously changes when plants are challenged by chewing insects and trigger both direct and indirect defenses through the HPL pathway (Bate and Rothstein, 1998; Blée, 1998; Croft *et al*., 1993; Wu and Baldwin, 2010). In functional terms, both AOS and HPL belong to the same superfamily of non-canonical cytochrome P450 enzymes but yet their amino acid sequences are distinct enough to permit their unique reaction mechanisms (Lee et al., 2008). While AOS interacts with LOX2 and AOC2, no interacting partners could be identified for HPL (Fig. 3). Last but not least, the chloroplast localization of AOS and HPL is quite distinct and helps to assure that AOS operates in the channeling of α-LeA to OPDA, whereas HPL drives volatile production.

In agreement with previous publications (Gilbert *et al*., 2008; Spivey and Ovádi, 1999), we conclude that compartmentalization of enzymes as well as organization into multi-protein complexes provides a highly specific cellular mechanism for controlling the flow of metabolites through key regulatory pathways and preventing unfavorable competing reactions. Work is in progress to obtain X-ray structural data for the identified LOX2-AOS-AOC2 complex from higher plant chloroplasts.

## EXPERIMENTAL PROCEDURES

### Plant growth

Wild-type *Arabidopsis*, *jaw-*D (Palatnik *et al*., 2003; Schommer *et al*., 2008), *aos* (Park *et al*., 2002; von Malek *et al*., 2002), *coi1* (Feys *et al*., 1994), *lox2* (Seltmann *et al*., 2010), *aoc2* (Stenzel *et al*., 2012) and *jar1* (Staswick *et al*., 1992) genotypes were used in this study. Homozygous *aos* plants were obtained by hand pollinating flowers with *aos* plants that had been generated by spraying flowering homozygous plants with 45 µM methyl jasmonate (MeJA). Seeds from an F_2_ population segregating for the *coi1* mutation were obtained as described (Feys *et al*., 1994). Plants were grown at 25 °C under standard conditions either under continuous white light illumination provided by fluorescent bulbs (30 W/m^2^) or 16 h light/8 h dark cycles.

### Generation of transgenic lines expressing AOS with COOH-terminally (His)_6_ or FLAG tags

Transgenic plants were generated expressing COOH-terminally (His)_6_ or FLAG-tagged AOS as described in the Supplemental Information section and used for affinity purification of proteins interacting with AOS *in planta*.

### Production of proteins and import into chloroplasts

Complementary DND (cDNA) clones for *AOS* (Laudert *et al*., 1996) and *HPL* (Froehlich *et al*., 2001) were cloned into appropriate vectors, allowing for their purification as (His)_6_- or Flag-tagged proteins. For routine chloroplast import assays, ^35^S-methionine- or ^14^C-leucine-labeled proteins were produced by coupled *in vitro* transcription/translation in wheat germ extracts. Radiolabeled proteins were added to 50 µL import assays consisting of 25 µL of doubly-concentrated import buffer, 10 µL of a plastid suspension containing 5 x 10^7^ *Arabidopsis* chloroplasts, and 2.5 mM Mg-ATP. All import reactions were performed at 23 °C for 15 min in darkness. Post-import protease treatment of plastids with thermolysin or trypsin and extraction of membranes with sodium carbonate, pH 11, or 1 M NaCl were carried out as described (Cline *et al*., 1984; Kessler and Blobel, 1996). Trypsin quenching was monitored by Western blotting using TIC110 as diagnostic marker and employing a trypsin inhibitor (Jackson *et al*., 1998). Plastid sub-fractionation into envelopes, stroma and thylakoids was made according to Li et al. (1991). Protein was extracted and precipitated with trichloroacetic acid (5% (w/v) final concentration), resolved by SDS-PAGE on 10-20 % (w/v) polyacrylamide gradients (Laemmli, 1970) and detected by autoradiography.

### Isolation of AOS-containing higher molecular mass complexes

AOS-(His)_6_ was imported into isolated *Arabidopsis* chloroplasts and AOS-(His)_6_ complexes released with detergent. For comparison, AOS-containing complexes were purified from plants expressing Flag-tagged AOS (AOS-Flag; see SI section). Both types of complexes, in turn, were subjected to size exclusion chromatography on Superose 6 (column model HR10/10, GE Healthcare) and individual fractions harvested and traced for the presence of AOS by Western blotting using AOS- or FLAG-specific antibodies (Laudert *et al*., 1996) and an enhanced chemiluminescence kit (ECL, GE Healthcare).

### Crosslinking

Crosslinking was carried out using ^125^I-*N*-[4[(p-azidosalicyl-amido)butyl]-3’(2-pyridyldithio) propionamid (^125^I-APDP)-derivatized precursors, essentially as previously described (Ma *et al*., 1996). Final protein samples were separated by reducing or non-reducing SDS-PAGE and ^125^I-labeled proteins detected by autoradiography.

### Enzyme activity measurements

Activity measurements using the isolated native and reconstituted complexes or free enzymes were carried out as described in *Supplemental Materials*. HPLC and GC-MS analyses used to identify and quantify substrates and products of the LOX2, AOS and AOC2 reactions were performed according to Holtman *et al*. (1997) and Zerbe *et al*. (2007). Capillary chiral GC analysis was used to demonstrate the optical purity of the *cis*-(+)-enantiomer (>95%) that was reconfirmed by TLC, HPLC and GC analyses with a synthetic standard (Zerbe *et al*., 2007).

### Yeast two-hybrid screens

Split-ubiquitin yeast two-hybrid screens for membrane bound proteins were carried out according to the manufacturer’s instructions using a commercial system (Dualsystems Biotech AG, Switzerland). Vector construction is described in *Supplemental Materials*.

### Bimolecular fluorescence complementation

The BiFC experiments were carried out according to (Walter *et al*., 2004). Vector construction and protoplast transformation is summarized in *Supplemental Materials*.

### Molecular modelling

Molecular modeling methods and tools are described in the *Supplemental Material* section.

### Miscellaneous

Western blotting was carried out according to (Towbin *et al*., 1979), using the indicated antisera and enhanced chemiluminescence (ECL, GE Healthcare) or antirabbit, anti-goat, alkaline phosphatase systems.

## ACKNOWLEDGMENTS

We are grateful to John Froehlich, Michigan State University, East Lansing, USA, Jörg Lehmann, formerly at Leibnitz Institute of Plant Biochemistry, Halle/Saale, Germany, Klaus Apel, formerly at Institute for Plant Sciences, ETH Zurich, Zurich, Switzerland, as well as Felix Kessler, Université Neuchatel, Neuchatel, Switzerland, and Danny J. Schnell, The University of Massachusetts, Amherst, USA, for gifts of cDNA clones and antibodies. The authors are also grateful to Nikolaus Amrhein, ETH Zurich, Zurich, Switzerland, for critically reviewing the manuscript and for his valuable comments. We thank Maik Hoffmann, Ruhr University Bochum, Germany, for technical assistance. This work was supported by a Marie-Curie grant of the European Communion [FP7-PEOPLE-CIG-2011-303744 to SP].

